# Comparative Single-Cell Profiling of CRISPR Knockout and Interference Defines Modality-Specific Strengths in Functional Genomics

**DOI:** 10.64898/2026.08.26.747303

**Authors:** Nikhil Gupta, Andy Sayer, Malwina Prater, Chara Mastrokalou, Khalid Saeed, Carlos Company, Curtis Hart, Ricardo Miragaia, Ashutosh Trehan, Functional Genomics Centre, Ultan McDermott, Magdalena E Strauss, Douglas Ross-Thriepland, David Walter, Alex Kalinka

## Abstract

Pooled CRISPR screens coupled with single-cell RNA sequencing enable high-throughput functional interrogation of gene regulatory networks, yet systematic comparisons of CRISPR knockout (CRISPRko) and CRISPR interference (CRISPRi) remain limited. We established a single-cell CRISPRko workflow and benchmarked it against CRISPRi using 87 sgRNAs targeting 29 unfolded protein response genes. CRISPRko generated transcriptional phenotypes highly concordant with CRISPRi, and induced comparable pathway-level responses albeit with distinct on-target and off-target effects between modalities. While CRISPRi allows direct assessment of target gene repression, CRISPRko provides an effective complementary approach for complete loss-of-function studies, expanding the toolkit for single-cell functional genomics.

## Background

Pooled CRISPR screening enables systematic, genome-scale identification of genes regulating diverse cellular processes, including therapeutic response, but conventional screens are largely limited to low-dimensional phenotypic readouts that do not capture transcriptional programs or cellular heterogeneity [1]. Coupling pooled CRISPR perturbations with single-cell RNA sequencing (scRNA-seq) overcomes these limitations by linking genetic perturbations to transcriptome-wide phenotypes in individual cells. Methods including Perturb-seq, CRISP-seq and CROP-seq established this approach, enabling the resolution of pathway activity, cellular state transitions and heterogeneous responses to genetic perturbation, with subsequent advances extending these methods to genome-scale studies [2-10].

Most single-cell CRISPR screens have employed CRISPR interference (CRISPRi), in which catalytically inactive Cas9 fused to transcriptional repressor domains silences gene expression without altering the coding sequence [11, 12]. CRISPRi provides efficient and reproducible gene repression and is particularly valuable for studying essential genes [12, 13]. In contrast, CRISPR knockout (CRISPRko) uses nuclease-active Cas9 to generate loss-of-function mutations through error-prone non-homologous end joining, enabling complete disruption of gene function but producing heterogeneous editing outcomes [13-15]. CRISPRko can additionally enable isoform-selective disruption by targeting unique coding exons, providing an approach to dissect isoform-specific functions that are generally inaccessible through promoter-directed transcriptional repression. Although CRISPRko is widely used in pooled functional genomic screens, its performance relative to CRISPRi in single-cell transcriptomic assays has not been systematically evaluated.

Here, we established a single-cell CRISPRko workflow and benchmarked it directly against CRISPRi using perturbations of the unfolded protein response (UPR) pathway [2, 8, 16]. We assessed the concordance of gene- and pathway-level transcriptional responses between the two modalities while evaluating their respective advantages and limitations for single-cell functional genomics.

## Results and Discussion

To establish a single-cell CRISPRko workflow and benchmark its performance against CRISPRi, we first optimised key experimental parameters, including guide capture strategy and library design. The unfolded protein response (UPR) pathway was selected as a benchmark system because it contains well-characterised transcriptional programmes spanning the three canonical branches of IRE1α, PERK, and ATF6 signalling [2, 8, 16, 17]. We designed an initial CRISPRko library targeting 29 UPR-associated genes using three sgRNAs per gene and evaluated both 3′ and 5′ single-cell capture strategies (Figure S1A). Both approaches generated comparable high-quality datasets following standard single-cell quality control filtering (Figure S1B, S1D), with similar pseudobulk transcriptional profiles between capture methods (Figure S1C). Comparison of perturbation signatures demonstrated strong concordance between 3′ and 5′ capture approaches, supporting the ability of both strategies to recover CRISPRko-associated transcriptional responses (Figure S1E). To further benchmark the ability of CRISPRko to generate transcriptional profiles comparable to established single-cell CRISPRi approaches, CRISPRko capture datasets were compared against the published Replogle et al. (2020) CRISPRi Perturb-seq dataset. Following equivalent quality control filtering (Figure S2A, S2B), the published CRISPRi dataset demonstrated expected target transcript repression (Figure S2C). CRISPRko perturbation signatures generated using both 3′ and 5′ capture showed concordant transcriptional responses with the CRISPRi Replogle dataset (Figure S2D, S2E), demonstrating the feasibility of single-cell CRISPRko as an alternative perturbation modality with similar transcriptional signatures.

As 5′ capture enables direct sgRNA detection using the native scaffold sequence without requiring additional vector modification, this approach was selected for subsequent experiments due to its broader compatibility. Initial evaluation of the pilot CRISPRko library identified variability in guide performance. To improve perturbation consistency, guides were refined using a combination of computational prediction and empirical performance metrics [18], generating a second-generation CRISPRko library (CRISPRko v2, see Methods).

For direct benchmarking, K-562 cells expressing Cas9 or dCas9-ZIM3 were transduced with matched CRISPRko v2 and CRISPRi sgRNA libraries targeting UPR pathway genes, respectively, and profiled after one week using whole-transcriptome sequencing, direct sgRNA capture, and targeted enrichment of genes associated with the UPR, DNA damage response, and apoptosis pathways (Figure 1A, S3A, S5A) [19, 20]. Cas9 and dCas9-ZIM3 activity was independently validated by targeting CD69 and measuring surface CD69 expression by flow cytometry, with both systems showing the expected guide-dependent changes in CD69 expression (Figure S3B). The sgRNAs targeting UPR-associated genes encompassed a broad spectrum of gene essentiality, with DepMap gene effect scores ranging from approximately 0 (non-essential) to −2 (strongly essential), to evaluate perturbation performance across genes with the full range of fitness levels (Figure S4A). Comparable numbers of high-quality cells were recovered from CRISPRko and CRISPRi datasets following filtering based on gene counts, transcript counts, and mitochondrial transcript content (Figure 1B, S3C). Similarly, targeted sequencing datasets showed comparable quality metrics and perturbation recovery between CRISPRi and CRISPRko conditions (Figure S5B, S5D). Global transcriptional profiles were highly similar between modalities, with comparable gene expression patterns and no evidence of broad transcriptional bias introduced by either perturbation strategy (Figure 1C, S5C).

**Figure 1.**
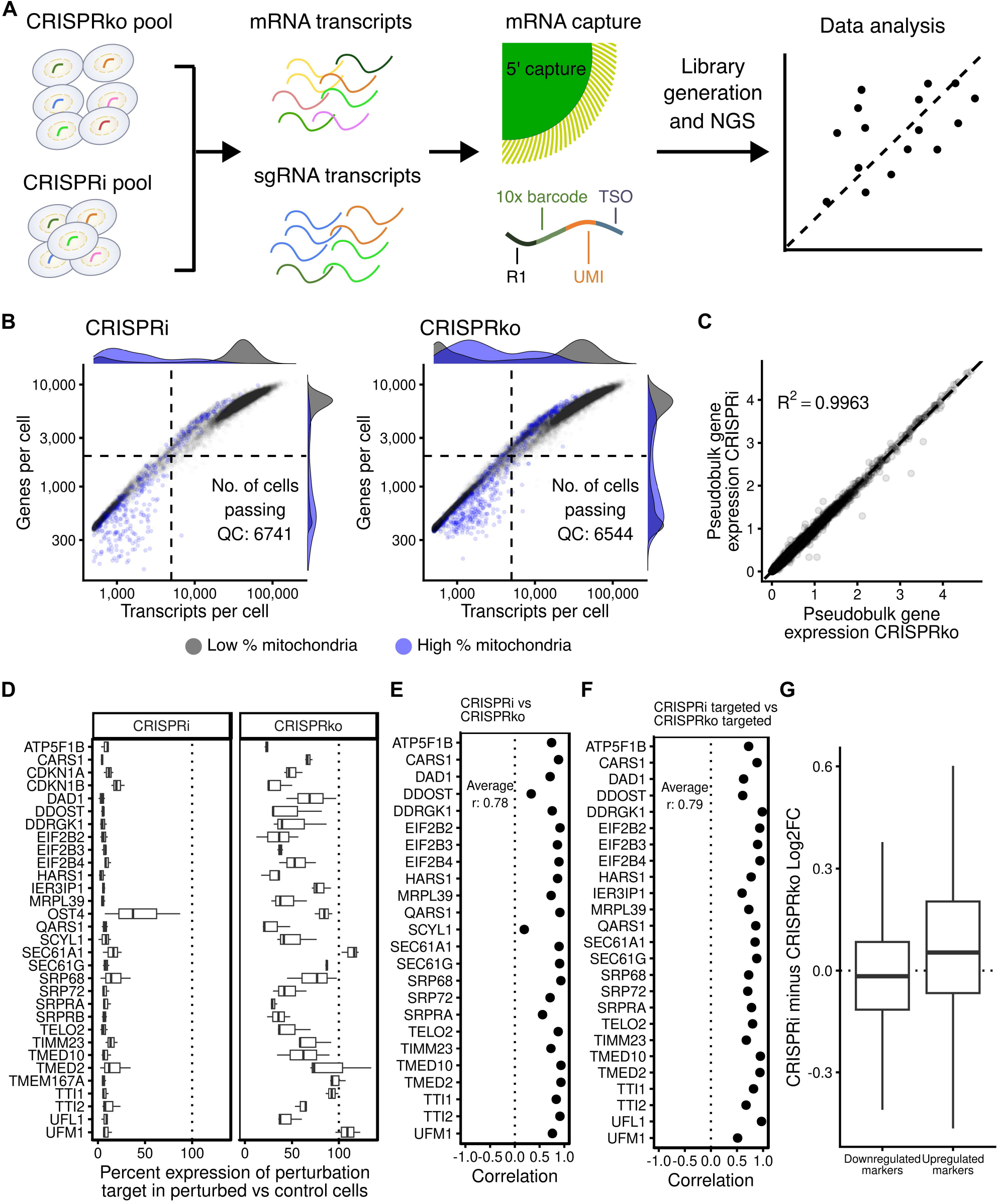
a. Schematic showing how CRISPRko and CRISPRi experiments were performed. b. Captured cells were filtered according to quality control criteria to ensure high quality cells were being assessed: genes per cell > 2000, transcripts per cell > 5000, mitochondria percentage < (Q3 + 3 IQR). Cells that do not meet the mitochondrial threshold are shown in blue; all other cells are shown in grey. Similar numbers of cells passing quality thresholds were obtained for CRISPRi and CRISPRko datasets. c. Pseudobulk gene expression for CRISPRi and CRISPRko shows averaged normalised gene expression remains similar between technologies. d. Percentage gene expression of perturbation-target in perturbed cells relative to expression of the same gene in non-targeting control (NTC) cells. Boxplots composed of three sgRNAs per perturbation target, for the CRISPRko and CRISPRi datasets. Clear target-gene repression observed with CRISPRi. e. CRISPRi vs CRISPRko marker gene correlation. For each perturbation, differentially expressed marker genes from either experiment are identified. The Pearson correlation of log2FC between datasets is calculated for each perturbation. Mean r across perturbations with >20 genes is indicated. f. CRISPRi targeted vs CRISPRko targeted marker gene correlation. For each perturbation, differentially expressed marker genes from either experiment are identified. The Pearson correlation of log2FC between datasets is calculated for each perturbation. Mean r across perturbations with >20 genes is indicated. g. Per-dataset and per-perturbation marker genes were identified. For each perturbation, marker genes that showed absolute shrunken log2FC > 0.1 and an adjusted p value of <0.05, and were found both in CRISPRi and CRISPRko, were taken forward. The difference in log2FC (CRISPRi minus CRISPRko) across all perturbations and common marker genes was calculated. Median value is indicated with thick horizontal black line. CRISPRi showed greater absolute log2FC (sign test: upregulated markers p < 1e-15; downregulated markers p < 1e-05) although the magnitude of this difference is small.

As expected from their distinct mechanisms, CRISPRi and CRISPRko differed in their ability to provide direct target transcript readouts. CRISPRi perturbations resulted in consistent reduction of target gene expression relative to non-targeting controls, reflecting transcriptional repression mediated by dCas9-ZIM3 (Figure 1D). By contrast, CRISPRko perturbations did not consistently reduce target transcript abundance, consistent with a DNA-level disruption mechanism in which frameshift or nonsense mutations do not necessarily alter transcript stability. Therefore, unlike with CRISPRi, target transcript abundance cannot be used as a direct per-cell indicator of CRISPRko perturbation efficiency.

Despite this mechanistic difference, CRISPRko and CRISPRi generated highly concordant transcriptional responses. Perturbation-associated marker genes identified independently in each dataset showed significant correlation in log2 fold changes between modalities (Figure 1E). One notable exception was SCYL1, where CRISPRi-mediated knockdown produced a distinct transcriptional profile from CRISPRko despite efficient target gene repression, suggesting that partial transcriptional inhibition did not fully recapitulate the effects of complete gene disruption (Figure S4B). Similar concordance was observed using targeted sequencing datasets designed to increase sensitivity for low-abundance transcripts, with targeted CRISPRi and CRISPRko profiles showing comparable marker gene correlations (Figure 1F). Across shared marker genes in all perturbations, CRISPRi produced larger absolute transcriptional changes than CRISPRko and larger on-target expression scores than CRISPRKo, although the magnitude of these differences was small (Figure 1G; Figure S7A). This trend was also observed across the number of differentially expressed genes detected per perturbation (Figure S4D, S4E).

To determine whether differences between CRISPRko and CRISPRi affected biological interpretation, we first compared pathway-level responses associated with UPR activation. Both CRISPRko and CRISPRi perturbations reproduced established UPR branch-specific transcriptional programmes corresponding to IRE1α, PERK, and ATF6 signalling (Figure 2A). Hierarchical clustering of scaled transcriptional responses demonstrated similar grouping of perturbations across CRISPRi, CRISPRko, and published Replogle datasets, with conserved pathway-specific signatures (Figure S6A, S6B, S6C). Second, gene set enrichment analysis further demonstrated comparable activation of stress-response programmes, including hallmark pathways associated with cellular stress, DNA damage response, and apoptosis (Figure 2B) [19, 20] demonstrating that CRISPRko and CRISPRi both capture biologically meaningful downstream consequences of gene disruption.

**Figure 2.**
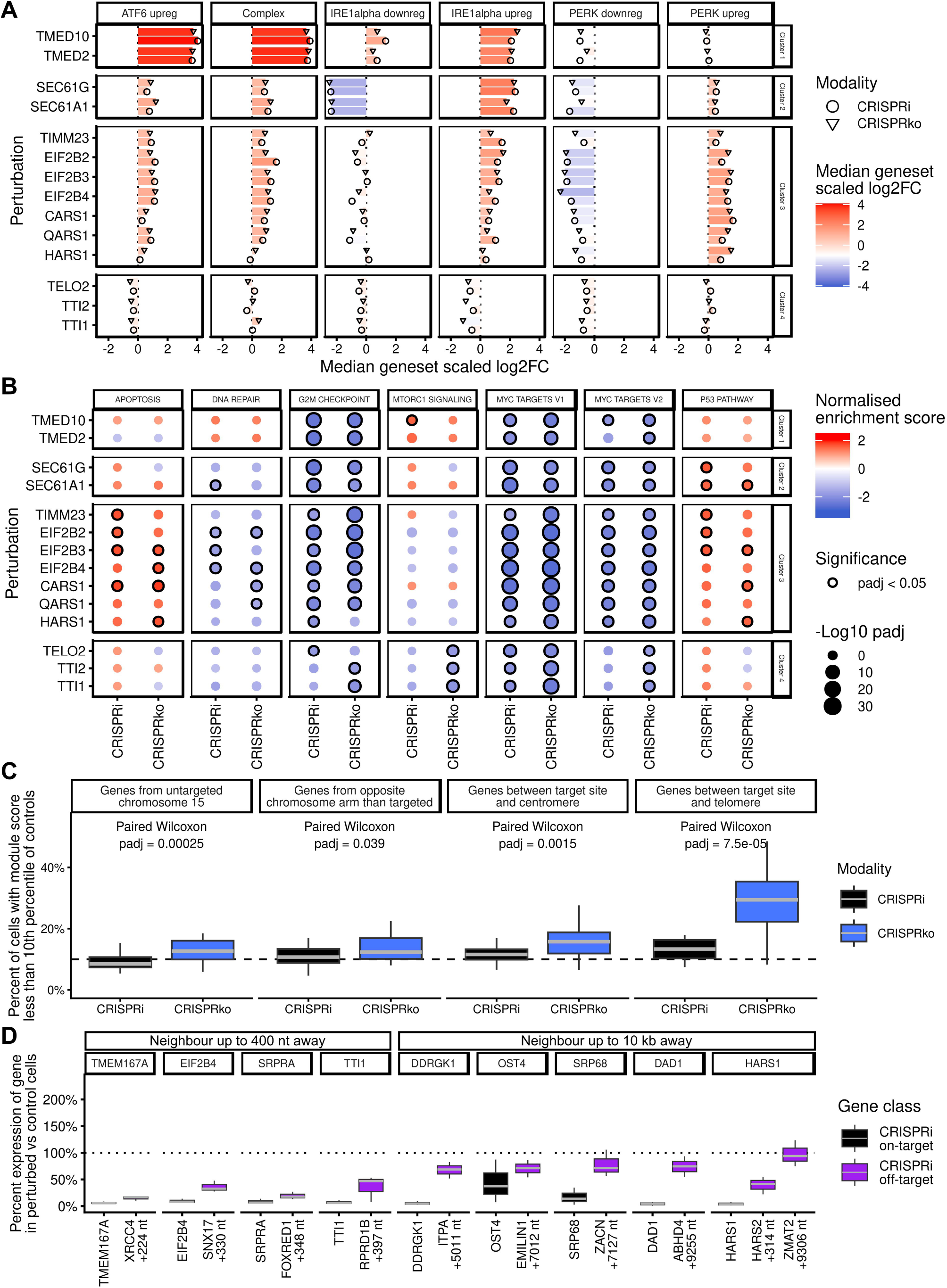
a. UPR pathway branch activation. CRISPRi and CRISPRko log2FC (gene expression in perturbed cells vs in control cells) was scaled by per-gene standard deviation in each dataset. Genes were grouped into genesets reflecting different UPR pathway branches described in Adamson et al., 2016. Median scaled log2FC is shown for each perturbation and geneset. CRISPRi shown with cricles; CRISPRko shown with triangles. b. Geneset enrichment analysis for selected HALLMARK genesets for CRISPRi and CRISPRko experiments. Shrunken Log2FC values were used as input for gene ranking. Normalised enrichment score is plotted and significance (padj < 0.05) is indicated with black outline. c. Chromosome region off-target effects were assessed by calculating module scores for different chromosome regions in perturbed and control cells (see methods). Cells were classified as showing low expression if their module score was below the 10th percentile of control cells, and their percentage per perturbation plotted. CRISPRko showed an increased percentage of cells with low expression in telomeric regions compared to CRISRPi p < 0.0001. d. Local off-target effects were assessed by identifying MANE Select transcripts with a transcriptional start site (TSS) within 10 kb of each CRISPRi target site. For perturbation-targets with a neighbouring TSS, the percentage gene expression of the perturbation target and neighbour in perturbed cells relative to expression of the same genes in non-targeting control (NTC) cells is shown for CRISPRi. The chromosomal distance between the furthest away CRISPRi sgRNA and the neighbouring gene TSS is indicated. CRISPRi was observed to repress the neighbouring genes at distances less than 400 nt.

The ability of CRISPRko to recover pathway-level phenotypes comparable to CRISPRi extends previous observations from bulk functional screens into the single-cell transcriptomic setting. Previous comparisons of CRISPRko, CRISPRi, and RNA interference approaches have demonstrated that optimised CRISPRko and CRISPRi libraries perform similarly for functional screening applications, despite differences in perturbation mechanism [13, 15]. Our findings demonstrate that CRISPRko and CRISPRi generate convergent biological information when coupled with single-cell transcriptomic readouts.

Finally, we assessed CRISPRi and CRISPRko off-target effects in terms of the transcriptional response at three scales: global, regional and local. CRISPRko was found to have an increased global off-target score (greater magnitude of expression change across the transcriptome after removing the on-target perturbation signature) (Figure S7B). Regionally, CRISPRko, showed a greater proportion of cells with low average expression of genes between the sgRNA target site and the telomere (Figure 2C). This effect was not observed in CRISPRi, nor was it seen in centromeric or opposite chromosome arm regions, suggesting it was CRISPRko and telomeric-region specific. We chose three perturbations to assess in more detail, which showed strong TSS-to-telomere repression and with a distance to the telomere greater than 50 Mb; we found that the CRISPRko-specific reduction of expression begins specifically at the sgRNA-target site and continues until the end of the chromosome (Figure S8A, S8B). The excess of CRISPRko cells showing repression in this region does not appear to be strongly related to the cell cycle phase (Figure S8C), distance between the sgRNA target site and the telomere (Figure S8D), nor the essentiality of the target perturbation (Figure S8E). We observed a weak association with the number of essential genes in the telomeric region (Figure S8F). However, with the current dataset, we do not fully understand this so it would require further investigation. Locally, we found that nine of our target-genes were within 10 kb of a neighbouring gene’s transcriptional start site (TSS). For both modalities, we assessed the repression of these target genes and their neighbours. With CRIPSRi, we observe strong repression of neighbouring genes within 400 nt of a CRISPRi sgRNA, and weak repression beyond 5 kb of a CRISPRi sgRNA (Figure 2D); effects not observed with CRISPRko (Figure S7C).

While CRISPRko and CRISPRi provide broadly comparable biological profiles, their distinct mechanisms confer specific experimental advantages and disadvantages. CRISPRi produces a stronger on-target transcriptomic signal, with less off-target behaviour both globally and in telomeric regions, than CRISPRko. However, CRISRPi induces strong repression of close neighbouring genes, which can potentially confound the assessment of particular genes. CRISPRi remains particularly valuable for interrogating genes where complete loss of function would compromise cellular survival. Thus, the choice of modality should depend at least on the genes being assessed. Comparison with DepMap gene effect scores demonstrated that the genes targeted in our experiment encompassed a range of essentiality states [21], including genes with strong fitness effects (Figure S4A). Perturbations targeting genes with stronger fitness effects showed reduced recovery in both CRISPRko and CRISPRi datasets (Figure S3C; S4B).

An advantage of CRISPRi is that target transcript expression provides an intrinsic molecular readout of perturbation efficacy. In the perturb-seq context, that allows direct assessment of repression efficiency and confidence in perturbation assignment without additional validation. CRISPRko efficiency cannot be inferred solely from transcript abundance and may require additional analyses to infer editing outcomes. However, CRISPRko does provide important advantages when complete gene loss is required. Moreover, CRISPRko targets coding regions rather than being restricted to transcription start site windows, thus accessing a larger sgRNA design space and use of extensively validated genome-wide knockout library designs [22, 23].

Together, these findings establish single-cell CRISPRko as a robust and informative complement to CRISPRi. Neither modality is universally superior; rather, the optimal strategy depends on the biological question being addressed. CRISPRi is advantageous when transcriptional repression, essential gene interrogation, targeting gene amplifications/mutated isoforms in cancer cell models or pharmacological modelling of partial inhibition is required, whereas CRISPRko provides a direct approach for studying complete loss-of-function phenotypes and exploiting broader targeting flexibility. As single-cell technologies continue to expand in scale and multimodal capability, integrating CRISPRko and CRISPRi

within complementary or sequential experimental designs will provide a more comprehensive framework for functional genomic discovery and genotype-to-phenotype mapping [5, 9].

## Conclusions

This study establishes single-cell CRISPRko as a robust and informative perturbation strategy that performs comparably to CRISPRi for single-cell transcriptomic profiling. Across sgRNAs targeting the UPR pathway, both modalities generated highly concordant gene- and pathway-level transcriptional signatures despite their distinct mechanisms of action and on-target and off-target effects. CRISPRi is advantageous when transcriptional repression, essential gene interrogation, studying mutated/amplified gene isoforms in cancer models or direct assessment of perturbation efficiency is required, whereas CRISPRko is better suited to modelling complete loss-of-function and offers greater flexibility for sgRNA design and library selection. Together, these findings demonstrate that CRISPRko and CRISPRi offer distinct advantages and provide a practical framework for selecting the most appropriate perturbation strategy in future single-cell functional genomics studies.

## Methods

### Cell Lines and Culture Conditions

All cell lines were obtained from the American Type Culture Collection (ATCC). Cell line identities were confirmed by short tandem repeat (STR) profiling, and all lines tested negative for mycoplasma contamination prior to use. Mycoplasma testing was performed routinely at approximately six-month intervals using either the PCR-based Universal Mycoplasma Detection Kit (ATCC) or the luminescence-based MycoAlert Detection Kit (Lonza). K-562 cells, derived from a chronic myelogenous leukaemia (CML) patient, were maintained in RPMI-1640 medium (25 mM HEPES, 2.0 g/L NaHCO3, 0.3 g/L L-glutamine) supplemented with 10% fetal bovine serum (FBS) and 2 mM GlutaMAX. HEK293T cells, used exclusively for lentiviral packaging, were cultured in Dulbecco’s Modified Eagle Medium (DMEM) supplemented with 10% FBS. All cell lines were maintained at 37°C in a humidified atmosphere containing 5% CO2.

### Generation of Stable Cas9 and dCas9-ZIM3 Cell Lines

To generate K-562 cell lines with stable expression of either nuclease-active Cas9 or the CRISPRi effector dCas9-ZIM3, cells were transduced with lentiviral particles derived from in-house constructs pFGCLV1-Cas9 and pFGCLV1-dCas9-ZIM3, respectively (synthesised by Genewiz) [24]. Transduction was performed in the presence of 8 µg/ml polybrene (Merck) for 24 hours, after which the medium was replaced with fresh complete RPMI. Blasticidin selection commenced 48 hours post-transduction and was continued until stable integrants were enriched. Cas9 nuclease editing efficiency was validated to exceed 85% using a dual fluorescent protein reporter system comprising in-house constructs pFGC-BFP/GFP-empty and pFGC-BFP/GFP-sgGFP. Additional validation was performed using lentiviral CRISPRko guide constructs LV_pFGCLV1_CD69_N1 and LV_pFGCLV1_CD69_N2, which target the CD69 cell surface marker, with editing efficiency assessed by flow cytometry using PE anti-human CD69 antibody (Biolegend #310906). Transcriptional repression activity of the dCas9-ZIM3 construct was validated using CRISPRi guide construct LV_pFGCLV1_CD69_I1 and LV_pFGCLV1_CD69_I2, also targeting CD69, with knockdown efficiency similarly confirmed by flow cytometry. For both modalities, debris was excluded followed by doublets and non-transduced cells as determined by BFP expression. The AAVS1 population across both modalities was used to inform the CD69-positive threshold. The guide sequences are listed in supplemental file 1.

### Lentiviral Production

Lentiviral particles were produced in HEK293T cells using a third-generation packaging system. Briefly, 20 million HEK293T cells were seeded into T175 flasks the day prior to transfection to achieve approximately 90% confluency at the time of transfection. A transfection mixture was prepared by combining 30 µg of the transfer vector, 25 µg of the packaging plasmid psPAX2 (Addgene #12260), and 10 µg of the envelope plasmid pMD2.G (Addgene #12259) in 3 ml Opti-MEM (Thermo Fisher Scientific). A total of 195 µl of X-tremeGENE HP DNA Transfection Reagent (Roche, #06 366 546 001) was added to the plasmid mixture, gently mixed, and incubated at room temperature for 20 minutes to allow transfection complex formation. The complexes were then added dropwise to the HEK293T cells. The following morning, the medium was replaced with 30 ml of fresh complete DMEM. Approximately 60 hours post-transfection, the virus-containing supernatant was harvested and clarified by filtration through a 0.45 µm low-protein-binding membrane. The filtered supernatant was aliquoted and stored at -80°C until use.

### sgRNA Expression Construct Design

For the 3′ gene expression capture workflow, sgRNA expression constructs were cloned into an in-house lentiviral backbone incorporating the 10x Genomics CS1 capture sequence at the 3′ position of the sgRNA scaffold [8], enabling direct sgRNA detection via the 3′ Feature Barcode capture chemistry. This backbone was used for the pilot CRISPRko v1 library. To enable a direct and minimally confounded comparison of 3′ and 5′ capture performance, the same pool of CRISPRko v1-transduced K-562 Cas9 cells was processed in parallel through both capture workflows, eliminating inter-sample biological variability as a confounding factor in the capture modality comparison. For the subsequent 5′ gene expression capture workflow, sgRNA expression constructs were cloned into a standard backbone lacking the CS1 sequence, as sgRNA detection in this modality exploits the constant region of the sgRNA scaffold as a native capture site and does not require an additional appended capture sequence. This standard backbone was used for cloning both the refined CRISPRko v2 library and the matched CRISPRi library employed in the final benchmarking experiments.

### sgRNA Library Design

The CRISPRko v1 library (used for 3’ capture and 5’ capture experiments) was built by filtering the Vienna library for genes expected to affect the UPR pathway when knocked out [18]. Three sgRNA sequences were chosen per gene ranked by the Auto-pick column. Five sgRNAs targeting the genomic safe harbour region AAVS1 were added, and five non-targeting sgRNAs from the Yusa library were added [25]. The final library contained 97 constructs.

The second iteration of the CRISRPko library (CRISPRko v2) (used for CRISPRko and CRISPRko targeted experiments) was also based on the Vienna library [18]. Compared to the CRISPRko v1 library, sgRNA sequences targeting SRPRA and four cell checkpoint genes CDKN2A, CDKN2B, CDKN1A and CDKN1B were added, and YIPF5 was removed. A number of sgRNAs from CRISPRko v1 were identified for replacement [26, 27]. Replacement sgRNA sequences were chosen from the Vienna library ranked by the Auto-pick column [28, 29]. Six sgRNAs targeting genomic safe harbour regions (AAVS1 and the Rogi1 region [30]) were added, and 6 non-targeting sgRNAs from the Yusa library were added [25]. The final library contained 111 constructs.

The CRISPRi library (used for CRISPRi and CRISPRi targeted experiments) was built starting with UPR pathway sgRNA sequences from Replogle et al 2022 (Supplementary Table 4), filtered for sgRNA IDs from Nuñez et al 2021 (Table S3), and combined with UPR pathway affecting sgRNA sequences from Replogle et al 2020 (Supplementary Table 2) [8, 9, 31]. Its composition was aligned with the CRISPRko v2 library; relative to the CRISPRko v1 library, sgRNA sequences targeting SRPRA and four cell checkpoint genes CDKN2A, CDKN2B, CDKN1A and CDKN1B were added, and YIPF5 was removed. sgRNAs containing restriction sites (BbsI, BsmBI, MluI and BamHI), targeting overlapping genomic coordinates, or of different lengths, were replaced with sgRNAs from Replogle et al 2022 that did not meet these criteria, prioritised by their ranking in Replogle et al 2022 and targeting a single genomic location (Guidescan version 2.0.0 using the GCF_000001405.40_GRCh38.p14 genome assembly) [8, 9, 32]. Six sgRNAs targeting genomic safe harbour regions (AAVS1 and the Rogi1 region [30]) were added, and 6 non-targeting sgRNAs from the Yusa library were added [25]. The final library contained 111 constructs.

### sgRNA Library Cloning

An oligonucleotide pool comprising 72-mer sequences was synthesised by Twist Bioscience. The library pool was amplified by PCR using KAPA HiFi HotStart ReadyMix (Roche, #KK2602) with 10 ng of template per reaction and 300 nM of forward primer (5’-TATATATCTTGTGGAAAGGACGAAACACCG-3’) and reverse primer (5’-GCTGTTTCCAGCATAGCTCTTAAAC-3’). The PCR programme consisted of an initial denaturation at 95°C for 3 minutes, followed by 10 cycles of 98°C for 20 seconds, 61°C for 15 seconds, and 72°C for 15 seconds, with a final extension at 72°C for 1 minute. Three independent PCR reactions were performed to ensure adequate library representation; products were pooled and purified using the QIAquick PCR Purification Kit (Qiagen, #28106).

The in-house lentiviral sgRNA expression vector pFGCLV3 was digested with Esp3I (NEB, #R0734S) and gel-purified to remove the stuffer fragment. Purified PCR amplicons were inserted into the linearised vector using the NEBuilder HiFi DNA Assembly Kit (NEB, #E2623S) at a 5:1 molar ratio of insert to vector, following the manufacturer’s recommendations. Three independent assembly reactions were performed, pooled, ethanol-precipitated, and resuspended in nuclease-free water. Transformation was carried out in triplicate using NEB 10-beta electrocompetent E. coli (NEB, #C3020K) according to the manufacturer’s instructions. Transformed bacteria were pooled and plated onto 225 mm × 225 mm LB agar plates supplemented with ampicillin and cultured overnight at 30°C to achieve a minimum representation of 500-fold coverage per sgRNA. Plasmid DNA was extracted using the NucleoSnap Plasmid Midi Kit (Macherey-Nagel, #740494.50). Library representation and sgRNA distribution were confirmed by next-generation sequencing (NGS) of the extracted plasmid pool prior to lentiviral production.

### CRISPRi and CRISPRko Perturbation Screen

For lentiviral transduction of the sgRNA library, 20 million K-562 cells expressing either Cas9 (CRISPRko) or dCas9-ZIM3 (CRISPRi) were seeded into triple-layer flasks and transduced with the benchmarking sgRNA library virus at a multiplicity of infection (MOI) targeting 20–30% transduction efficiency, in the presence of 8 µg/ml polybrene (Merck). This low MOI ensured that the majority of transduced cells received a single lentiviral integrant, minimising the risk of multiple guide co-integration. The virus-containing medium was replaced with fresh complete medium the following day. Two days post-transduction, the proportion of BFP-positive cells, expressed from the library vector, was quantified by flow cytometry to confirm transduction efficiency within the target range of 20–30%. Puromycin selection commenced two days post-transduction and was continued until the BFP-positive fraction exceeded 90%, confirming enrichment of transduced cells. Throughout the selection and expansion period, cell cultures were maintained at sufficient scale to ensure a minimum representation of 500 cells per sgRNA at all timepoints, preserving library complexity. Cells were harvested for single-cell capture one week post-transduction, at which point guide-mediated perturbation was considered to be at steady state for both CRISPRko and CRISPRi modalities.

### Single-Cell Library Preparation

Single-cell transcriptomic libraries were generated using three complementary 10x Genomics Chromium-based workflows: 5′ gene expression with CRISPR screening, 3′ gene expression with CRISPR screening, and targeted gene expression. All libraries were prepared on the 10x Genomics Chromium platform.

### 5′ Gene Expression and CRISPR Screening Library Construction

For 5′ single-cell capture, single-cell suspensions were prepared at a concentration of 500 cells/µl and loaded at a volume of 33 µl per lane, yielding approximately 16,500 cells per lane with a target recovery of approximately 10,000 cells. Two lanes were used per sample, providing a combined target recovery of approximately 20,000 cells per sample. Cells were processed using the Chromium Next GEM Single Cell 5′ Reagent Kit v2 (Dual Index; PN 1000263/1000265) in conjunction with the 5′ CRISPR Kit (PN 1000451), the Library Construction Kit (PN 1000190), and the Chromium Next GEM Chip K Single Cell Kit (PN 1000286/1000287), according to the manufacturer’s protocol (CG000510, Rev B). Both gene expression and CRISPR feature barcode libraries were quality-assessed by capillary electrophoresis using an Agilent TapeStation and quantified prior to pooling and sequencing. Gene expression libraries were sequenced at a depth of 60,000 reads per cell and CRISPR feature barcode libraries at 5,000 reads per cell.

### 3′ Gene Expression and CRISPR Screening Library Construction

For 3′ single-cell capture, single-cell suspensions were prepared at a concentration of 400 cells/µl and loaded at a volume of 42 µl per lane, yielding approximately 16,800 cells per lane with a target recovery of approximately 10,000 cells. Two lanes were used per sample, providing a combined target recovery of approximately 20,000 cells per sample. Cells were processed using the Chromium Next GEM Single Cell 3′ Reagent Kit v3.1 (Dual Index; PN 1000268/1000269), the 3′ Feature Barcode Kit (PN 1000262), and the Chromium Next GEM Chip G Single Cell Kit (PN 1000120/1000127), following the manufacturer’s protocol (CG000316, Rev D). Post-construction quality control was performed and libraries were quantified prior to sequencing. Gene expression libraries were sequenced at a target depth of 60,000 reads per cell and CRISPR feature barcode libraries at 5,000 reads per cell.

### Targeted Gene Expression Library Construction

Targeted gene expression profiling was performed exclusively for the CRISPRko versus CRISPRi benchmarking experiment as a downstream hybridisation capture enrichment step, using the 10x Genomics Targeted Gene Expression workflow per the manufacturer’s protocol (CG000293, Rev G). The targeted libraries were derived from the same cDNA pools generated during the standard 5′ gene expression capture workflow, ensuring that the whole transcriptome and target enriched libraries originate from identical biological samples, enabling direct comparison between the two sequencing modalities. Two capture panel configurations were employed in parallel: the 10x Genomics Human Gene Signature Panel (16 rxns; PN 1000245) and a bespoke fully custom hybridisation panel designed and synthesised by Integrated DNA Technologies. The custom IDT panel was spiked into the hybridisation reaction alongside the pre-designed panel to enable simultaneous enrichment of both curated gene signatures and study-specific targets of interest. Libraries were prepared following the manufacturer’s protocol (CG000296, Rev B). Final targeted libraries were quality-assessed by capillary electrophoresis using an Agilent TapeStation and quantified prior to sequencing. Targeted enriched libraries were sequenced at a depth of 15,000 reads per cell.

### Sequencing

All libraries including gene expression, CRISPR feature barcode, and targeted enrichment libraries were sequenced on an Illumina NovaSeq 6000 instrument using S1 and S4 flow cells, with libraries assigned to flow cell format based on sample throughput and lane requirements. Read configuration and sequencing parameters followed 10x Genomics recommendations for each respective library type.

### Single-cell analysis and quality control

Gene expression quantification was performed with 10x Genomics Cell Ranger v7.1.0 and the GRCh38-2020-A reference annotation for all experiments [33]. For targeted sequencing experiments, 10x Genomics Cell Ranger v7.1.0 was used with the --target-panel argument to specify the custom set of probes.

Single cell RNA sequencing analysis was performed with Seurat v5.5.0 and R v4.5.3 [34]. Initial cell numbers were: CRISPRi 3’ capture 17725, CRISPRi 5’ capture 18761, CRISPRi 18343, CRISPRko 26898, CRISPRi targeted 13407 and CRISPRko targeted 14686. Cell level filtering was applied to exclude cells that had UMI counts <5000, unique genes <2000 and mitochondrial percentage > Q3 + 3×IQR. For targeted sequencing, cells were excluded if they had UMI counts <3000 and genes <700. Gene level filtering was applied to retain genes that showed expression in at least 10 cells.

10x Genomics Cell Ranger was used for initial sgRNA assignment and quantification with the pattern ‘GCTGTTTCCAGCATAGCTCTTAAAC(BC)’ for 5’ capture or ‘(BC)GTTTAAGAGCTATGCTGGAAACAGC’ for 3’ capture, wherein (BC) is the reverse complement of the sgRNA sequence for 5’ capture or the sgRNA sequence for 3’ capture. High confidence sgRNA assignments were achieved as follows: the log-transformed UMI counts for sgRNAs were fitted to a three-component skew-normal mixture model, with the cluster with the lowest mean corresponding to background noise, i.e. sgRNAs with low UMI counts were classified as background noise rather than being used for sgRNA assignment. A lower and upper UMI count threshold were derived from probabilities of belonging to this background noise component: the lower threshold corresponded to high background noise probability (>0.9), and the upper threshold to low background noise probability (<0.1). Cells were assigned to an sgRNA if: (1) that was the only sgRNA with UMI counts above the upper threshold; and (2) no other sgRNAs with UMI counts between the lower and upper thresholds were assigned to the cell.

Only cells with a single non-background noise sgRNA assigned to them were used in the final analysis. In addition, cell doublets were predicted with scDblFinder v1.24.10 and only cells predicted to be singlets were taken forward to final analysis [35]. Final cell numbers were: CRISPRi 3’ capture 7621, CRISPRi 5’ capture 8157, CRISPRi 6741, CRISPRko 6544, CRISPRi targeted 6695 and CRISPRko targeted 5627.

### Differential expression testing

DElegate v1.2.1 was used to identify differentially expressed genes between each perturbation and control perturbations (non-targeting controls), at both the sgRNA level and sgRNA-target level [36]. Firstly, DElegate was modified to avoid using the deprecated “slot” argument in Seurat function GetAssayData and use the “layer” argument instead. DElegate splits cells into three pseudoreplicates per perturbation for differential expression testing with DESeq2 v1.50.2 [37]. Log2 fold changes and shrunken log2 fold changes (‘ashr’ method) [38] were calculated, as were p-values and adjusted p-values. Adjusted p-values were further adjusted by multiplying by the number of perturbations being assessed. ‘Marker genes’ were called if they showed an absolute log2 fold change > 0.1 and an adjusted p-value < 0.05.

### Inter-modality marker gene correlation

To compare the effect of CRISPRi and CRISPRko on the wider transcriptome, log2 fold changes at the sgrna-target level were calculated in the manner described above. Per perturbation, genes were taken forward if they were found in both datasets, and if they were identified as marker genes (based on shrunken log2 fold change values) in either dataset. Per perturbation, the log2 fold change of these genes were extracted and used to calculate the Pearson correlation between CRISPRi and CRISPRko.

### UPR pathway branch activation

UPR pathway branch activation was determined with reference to gene sets representative of different UPR pathway branches, as described in Figure 3F of Adamson et al 2016 [2]. For each experiment and modality, the log2 fold change of each of these UPR branch genes was scaled by its standard deviation across all perturbations. Perturbations were hierarchically clustered and a consensus set of four clusters was derived after manual inspection across experiments.

### Gene set enrichment analysis

Gene set enrichment analysis was performed with the package fgsea v1.36.2 [39]. Hallmark gene sets were obtained for specific pathways and intersected with the genes present in the data [40]. Shrunken log2 fold changes were used to rank genes. Fgsea was run with 1000 permutations. Normalised enrichment score (enrichment score normalised to mean enrichment of random sample) and adjusted p-values were plotted.

### Global on- and off-target analysis

The on-target perturbation score was calculated by merging CRISPRi and CRISPRko data using Seurat v5.5 and performing PCA across all cells. Seurat’s CalcPerturbSig was used to identify the 20 closest modality-matched control cells in the common PCA space to each perturbation-target, and subtract their average gene expression from the gene expression matrix. For each modality, variable genes were identified by DElegate v1.2.1, as outlined above, filtered for padj < 0.05 and an absolute shrunken log2FC values > 0.25. For each perturbation-target, an intersection of variable genes across modalities was determined. At least 5 genes were required to proceed with calculating an on-target score. The average control-subtracted expression of genes from all control cells (combining both modalities) was subtracted from each cell, resulting in a control-centred matrix. For each perturbation target, every variable gene’s mean control-centred value was calculated, and divided by the vector’s magnitude to yield a ‘perturbation signature’ unit vector. For every cell of each perturbation, the control-centred vector of variable genes was projected onto the associated perturbation signature unit vector to yield an ‘on-target score’, describing the degree of alignment of the cell with the perturbation signature.

The total magnitude of the control-centred expression across all genes was calculated for every cell. This represents the total expression difference from control cells. The ‘off-target score’ was calculated as the orthogonal residual between the total magnitude of control-centred expression and the on-target score. The median off-target score of the control perturbations was subtracted from all off-target scores to centre control off-target values to zero.

Mixscale v.0.3.0 was used to generate a ‘mixscale score’, which has similarities to the ‘on-target score’ described above. Variable genes were identified between every perturbation-target and control, combining cells across modalities to ensure the variable genes are not modality-specific. Variable genes with a logFC > 0.1 were retained and at least 5 were required to generate a perturbation signature. The control-centred expression of the retained variable genes was used to generate a ‘perturbation signature’ unit vector. Other cells’ control-centered expression was projected onto this vector to return a perturbation score, which was then standardised by the control mean and standard deviation to yield a ‘mixscale score’.

### Regional off-target analysis

Regional off-target effects were assessed by calculating module scores (normalised average gene expression scores) for different gene sets using Seurat v5.5 AddModuleScore function. Module scores were calculated by identifying the sets of MANE-select v1.5 genes between each perturbation-target’s TSS and the (1) telomere; (2) centromere; (3) opposite chromosome arm; and (4) for an untargeted chromosome (chromosome 15). At least 17 genes were required to calculate a module score. For each perturbation, the module score was calculated for all cells of the perturbation and all control cells. The 10^th^ percentile module score from the control cells was used as a threshold to compare each perturbation’s module scores against. For each perturbation, the percentage of perturbed cells with a lower module score than the control cells’ 10^th^ percentile module score was calculated. For perturbations behaving similarly to controls this percentage was expected to be approximately 10 percent.

Regional off-target effects were also assessed by moving average of control-centred gene expression. MANE-select v1.5 genes were identified and ordered along each chromosome by position. For each perturbation, the genes located on the target gene’s chromosome were identified and a moving average of control-centred gene expression (expression of gene in perturbed cell minus the average expression of the same gene in control cells) was performed with a window of 51 genes along the chromosome. The median position of the 51 genes was associated with each moving average value. This score was used as the heatmap fill value (Figure S8A, S8B).

Regional off-target effects were compared to cell phase. For each modality, and each class of module score (‘Low’ or ‘Normal’), all cells of a perturbation were assigned to a cell state (G1, G2M or S) using Seurat v5.5.0 function CellCycleScoring; this was then converted to a percentage with respect to the total cells found in the group.

Regional off-target effects were further assessed by determining (per perturbation) the excess percentage of cells with a module score below the 10th percentile of controls in CRISPRko vs CRISPRi. This represents the CRISPRko-specific change in low module score, accounting for the effect of the perturbation. This was compared to: the distance between the target TSS and the telomere; the DepMap 26Q1 gene effect of the target in K-562 cells; and the number of genes in the TSS-to-telomeric region with a DepMap 26Q1 gene effect less than -1.

### Local off-target analysis

Local off-target effects were assessed by first identifying MANE-Select v1.5 genes with a TSS within 10 kb of each CRISPRi sgRNA target site. For the nine perturbation targets with a neighbouring TSS, the percentage gene expression of each target-gene and neighbouring gene relative to the same target-gene and neighbouring gene in control cells was calculated.

### Availability of data and materials

Sequencing data are available through ArrayExpress accession E-MTAB-17512, E-MTAB-17513, E-MTAB-17514, E-MTAB-17515, E-MTAB-17516, and E-MTAB-17518. Primer and guide sequences used in this study are listed in Supplementary File 1.

## Author contributions

NG and DW conceived the project. NG, KS and CH planned and performed the experiments. AS, MP, MES, CM, RM and CC performed the bioinformatics analysis. NG, CH, MES and AS wrote the manuscript. NG, AK, UM, MES, AT, DW and DRT supervised the project. All authors reviewed and approved the manuscript.

## Acknowledgements

We are very grateful to Marica Gaspari and to our colleagues at the Functional Genomics Centre for their contributions and useful advice. The authors gratefully acknowledge the Cambridge Stem Cell Institute (CSCI) Genomics Facility and Cancer Research UK Cambridge Institute (CRUK-CI) Genomics Facility for their support and assistance in this work. M.E.S. was supported by the Wellcome Trust (220442/Z/20/Z).

