## Supplemental Figures for "Comparative Single-Cell Profiling of CRISPR Knockout and Interference Defines Modality-Specific Strengths in Functional Genomics"

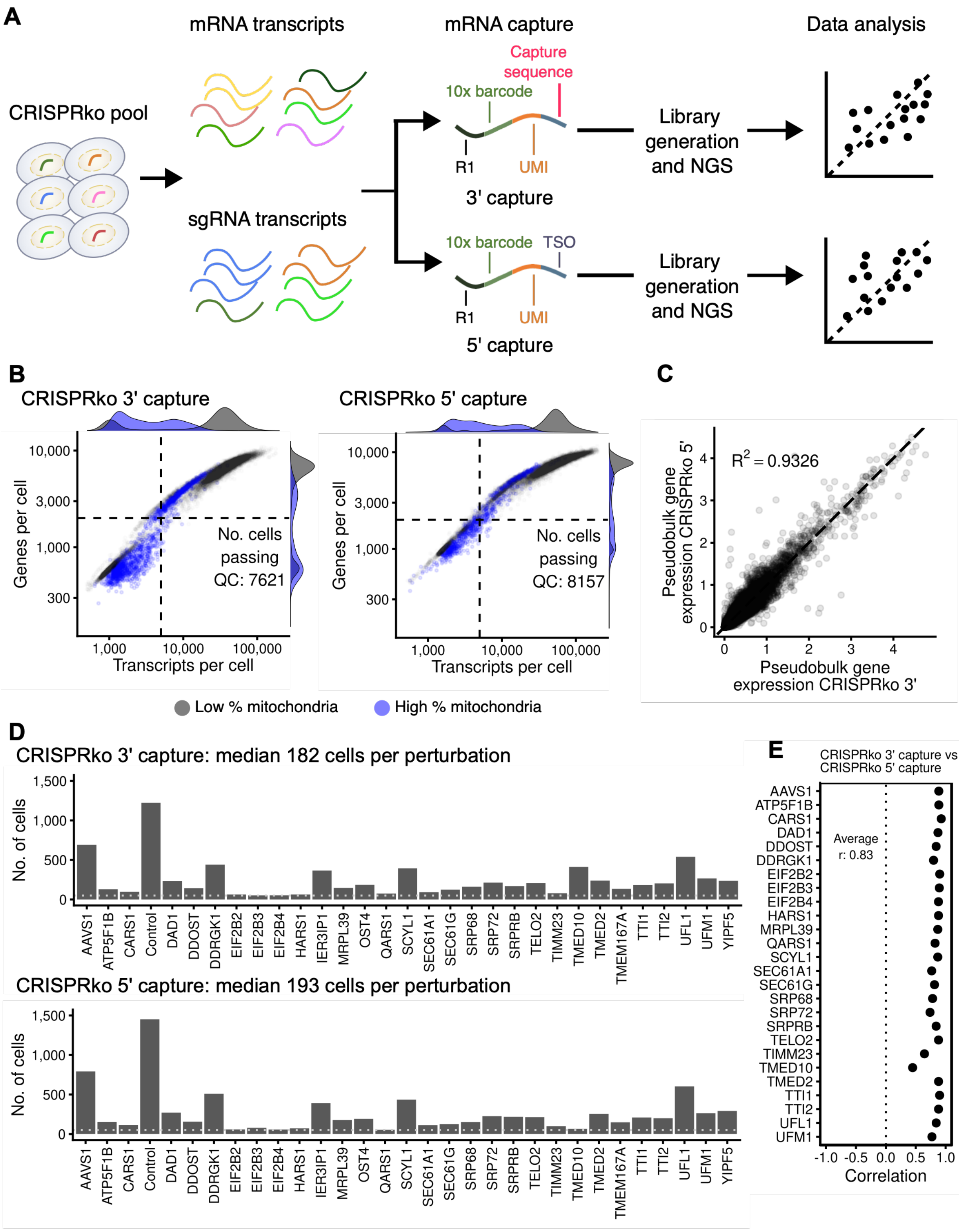


**Supplemental Figure 1.** a. Schematic showing how CRISPRko 3’ capture and 5’ capture experiments were performed. b. Captured cells were filtered according to quality control criteria to ensure high quality cells were being assessed: genes per cell > 2000, transcripts per cell > 5000, mitochondria percentage < (Q3 + 3 IQR). Cells that do not meet the mitochondrial threshold are shown in blue; all other cells are shown in grey. Similar numbers of cells passing quality thresholds were obtained for CRISPRi and CRISPRko datasets. c. Pseudobulk gene expression for CRISPRko 3’ capture and 5’ capture shows averaged normalised gene expression remains similar between technologies. d.

Number of cells that passed QC filtering per sgRNA target for CRISPRko 3' capture and 5' capture. Horizontal line indicates 50 cells. e. CRISPRko 3' capture vs CRISPRko 5' capture marker gene correlation. For each perturbation, differentially expressed marker genes from either experiment are identified. The Pearson correlation of log2FC between datasets is calculated for each perturbation. Mean r across perturbations with >20 genes is indicated.

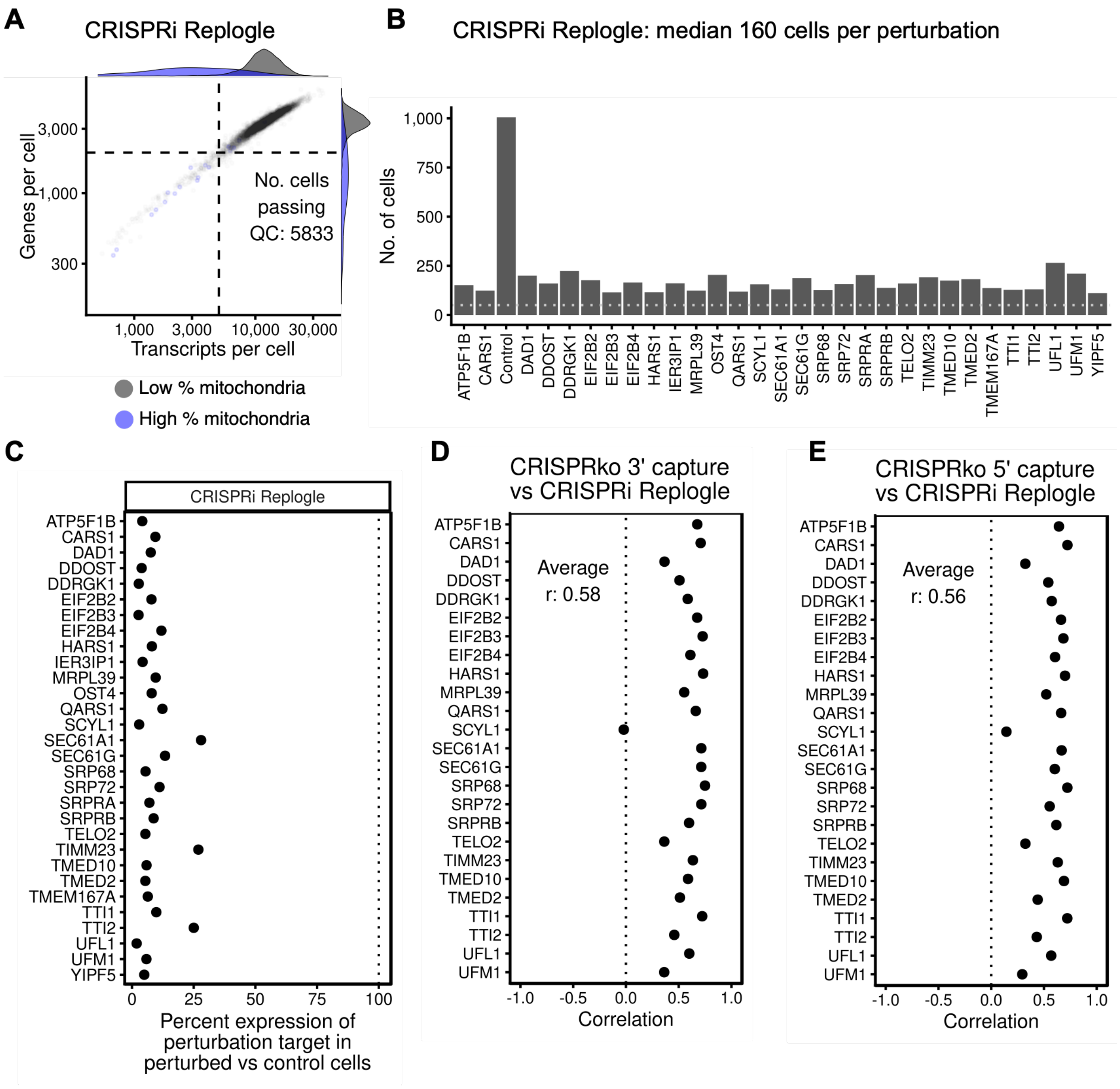


**Supplemental Figure 2.** a. Replogle et al., 2020 CRISPRi dataset: cells were filtered according to quality control criteria to ensure high quality cells were being assessed: genes per cell > 2000, transcripts per cell > 5000, mitochondria percentage < (Q3 + 3 IQR). Cells that do not meet the mitochondrial threshold are shown in blue; all other cells are shown in grey. b. CRISPRi Replogle dataset: number of cells that passed QC filtering per sgRNA target. Horizontal line indicates 50 cells. c. CRISPRi Replogle dataset: percentage gene expression of perturbation-target in perturbed cells relative to expression of the same gene in non-targeting control (NTC) cells. One sgRNA per gene. Clear target-gene repression observed. d and e. CRISPRko 3' capture (d) or CRISPRko 5' capture (e) vs CRISPRi Replogle marker gene correlation. For each perturbation, differentially expressed marker genes from either experiment are identified. The Pearson correlation of log2FC between datasets is calculated for each perturbation. Mean r across perturbations with >20 genes is indicated.

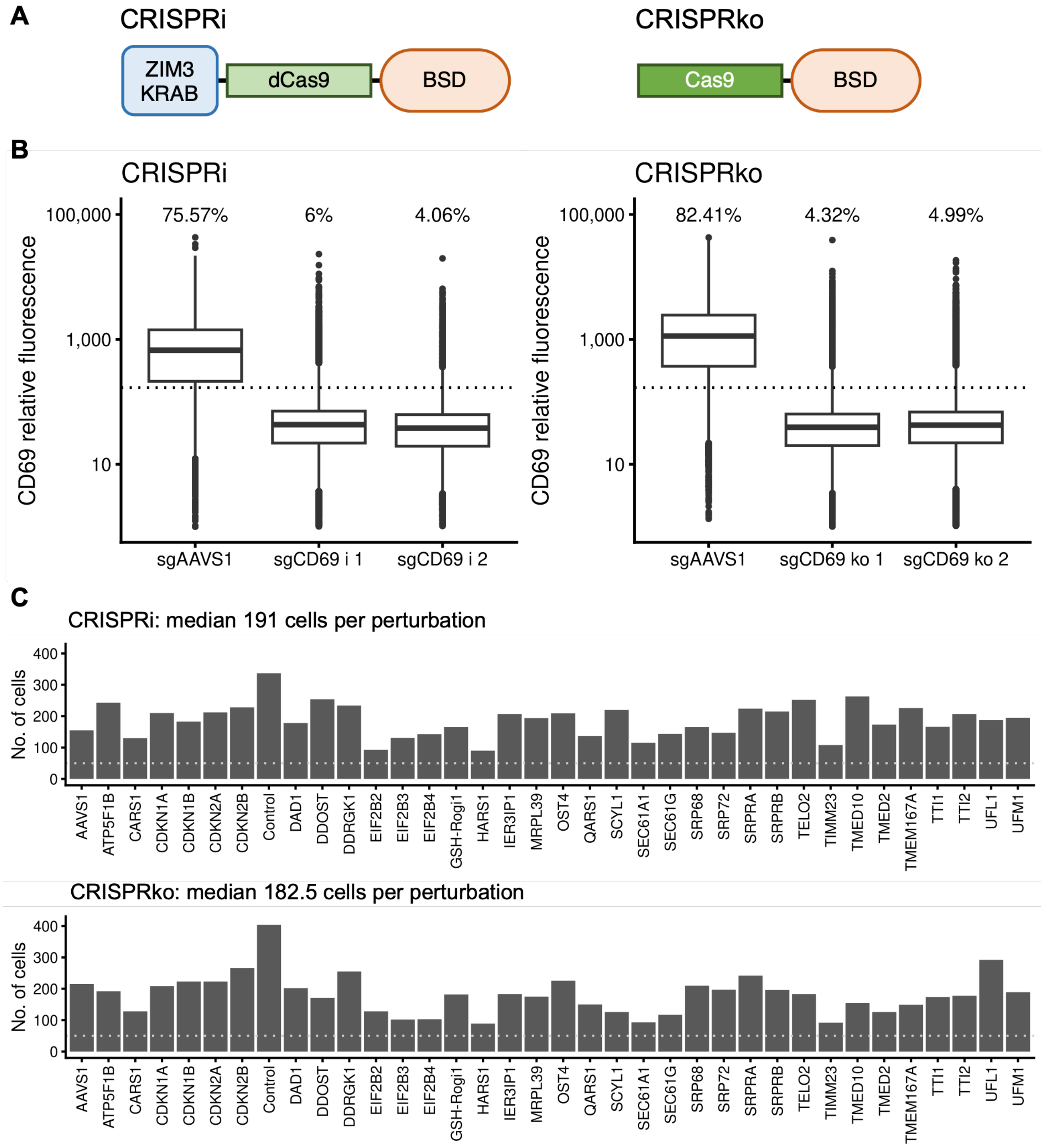


**Supplemental Figure 3.** a. Schematic showing the different constructs used for CRISPRi and CRISPRko experiments. b. Relative fluorescence of PE anti-human CD69 antibody in K-562 cells with dCas9-Zim3 (CRISPRi) and Cas9 (CRISPRko) for different sgRNAs targeting a neutral locus (AAVS1) or CD69. Percentages indicate the percent of filtered cells above the CD69 relative fluorescence gate (dotted line). c. Number of cells that passed QC filtering per sgRNA target for CRISPRi and CRISPRko. Horizontal line indicates 50 cells.

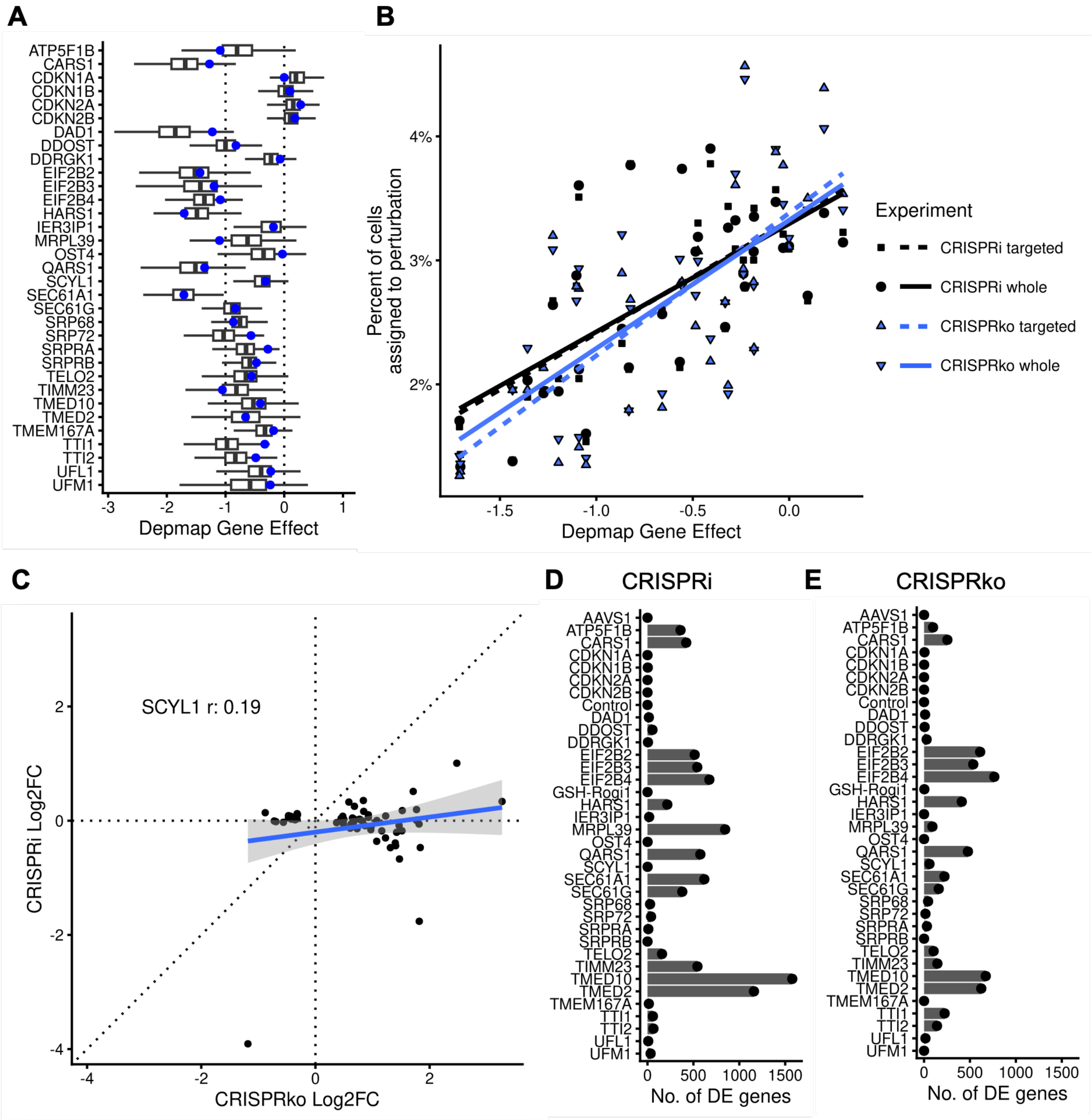


**Supplemental Figure 4.** a. DepMap 26Q1 gene effect score for perturbation targets in CRISPRi and CRISPRko libraries. Dotted line at 0 indicates non-essentiality and -1 indicates median of common essentials. Blue dots indicate Depmap Gene Effect of K-562 cells (ACH-000551). b. The perturbation targets in our screen were used to extract DepMap 26Q1 gene effect in K-562 (ACH-000551) cells. For each experiment, this was compared to the percentage of all cells passing QC that were assigned to each of the perturbation targets. A positive correlation is observed with no meaningful difference between CRISPR modality or gene expression sequencing method. c. Marker genes showing differential expression between SCYL1 perturbed cells and non-targeting control cells in CRISPRi and CRISPRko experiments were identified. Union of CRISPRi and CRISPRko marker genes was taken, and the log2 fold change (expression of gene in SYCL1 perturbed cells vs non-targeting control cells) of each gene is plotted in CRISPRi and CRISPRko experiments. d. and e. Marker genes are filtered to the set of genes for each perturbation with significant differential expression (padj < 0.05 and abs(log2FC) > 0.1). Total number of differentially expressed genes are plotted for CRISPRi and CRISPRko.

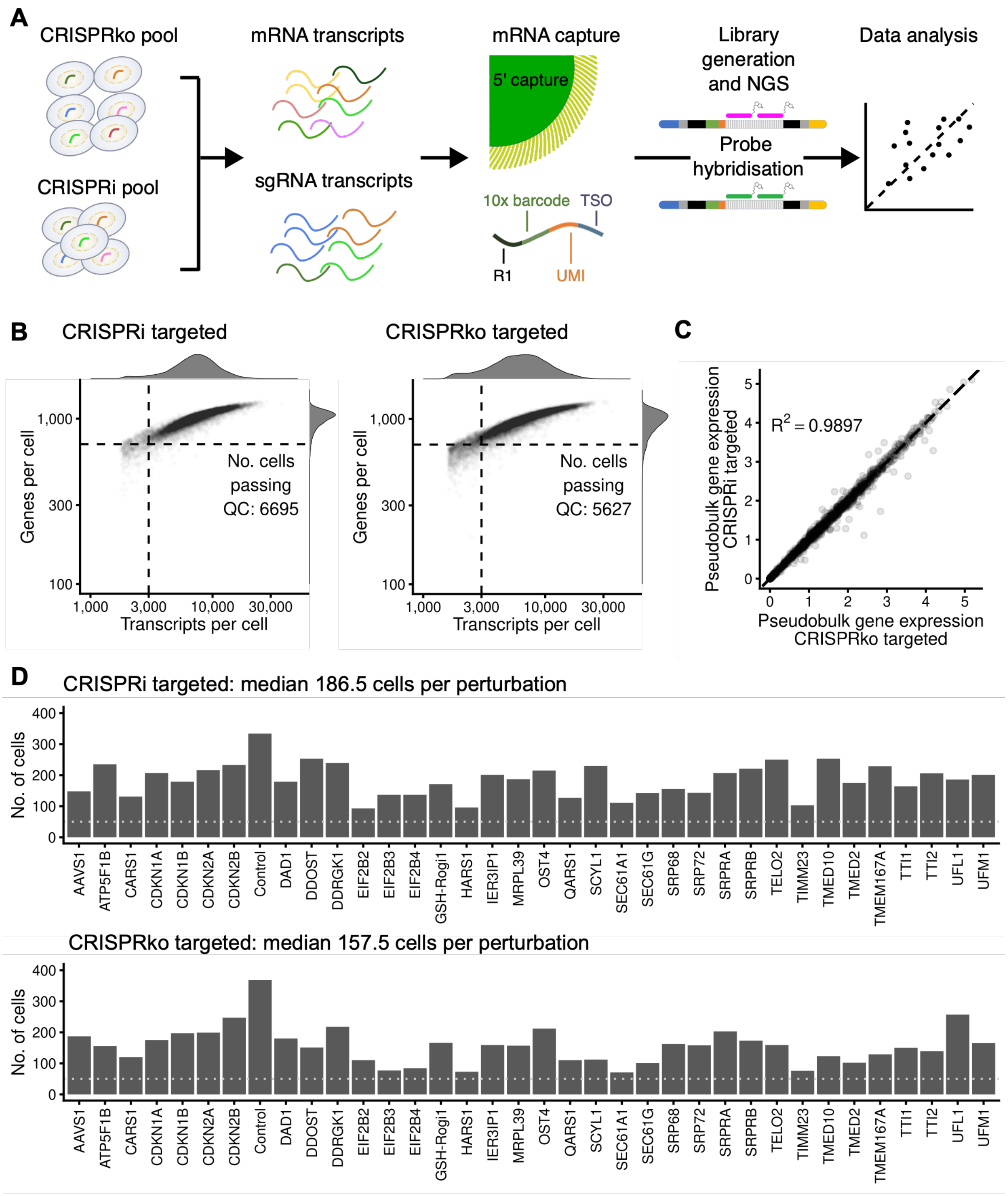


**Supplemental Figure 5.** a. Schematic showing how CRISPRi targeted and CRISPRko targeted sequencing experiments were performed. b. CRISPRi targeted and CRISPRko targeted captured cells were filtered according to quality control criteria to ensure high quality cells were being assessed: genes per cell > 700 and transcripts per cell > 3000. No mitochondrial filtering was performed. c. Pseudobulk gene expression for CRISPRi targeted and CRISPRko targeted shows averaged normalised gene expression remains similar between technologies. d. Number of cells that passed QC filtering per sgRNA target for CRISPRi targeted and CRISPRko targeted. Horizontal line indicates 50 cells.

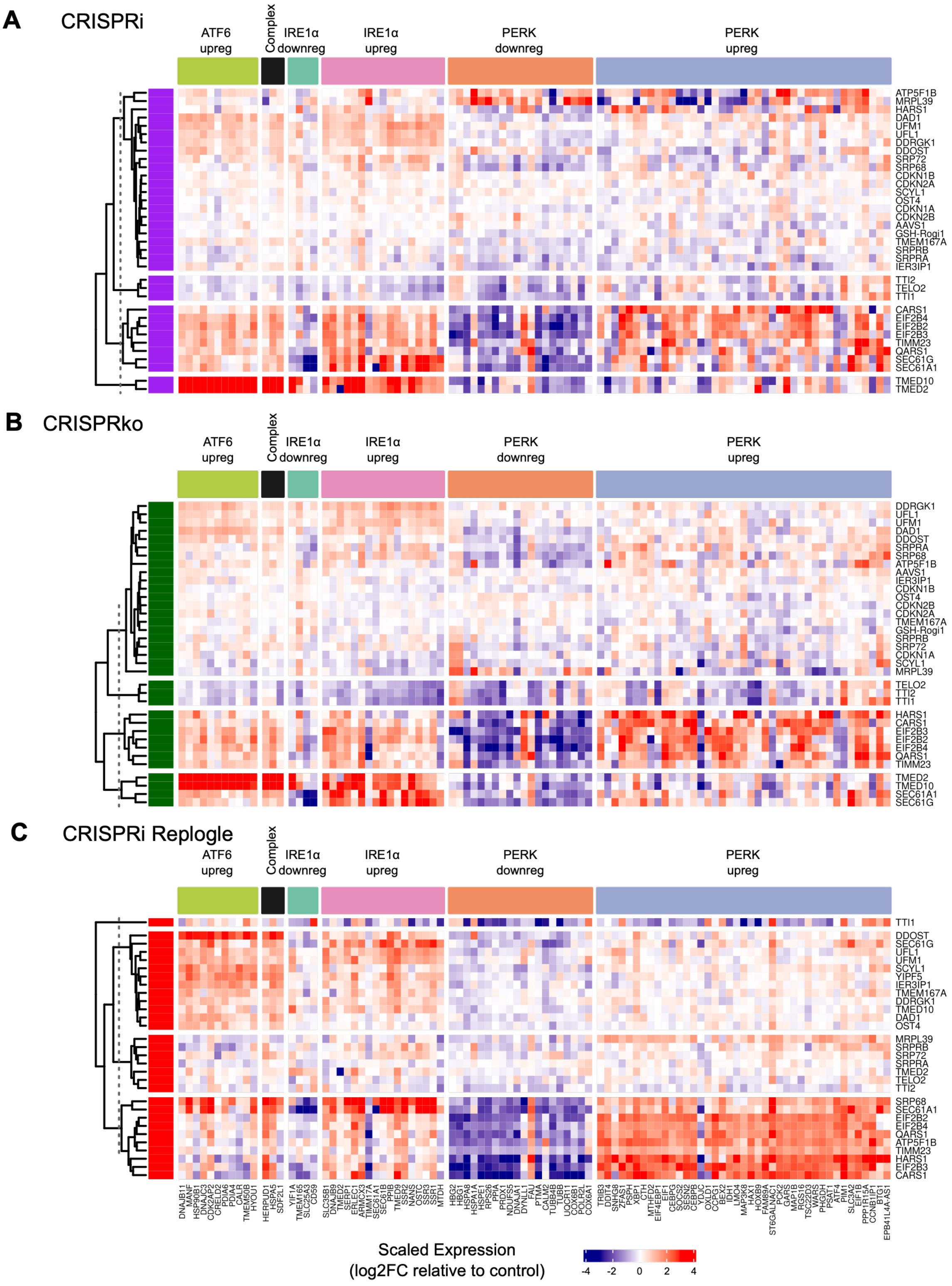


**Supplemental Figure 6.** Scaled log2FC heatmap for each of the (a) CRISPRi, (b) CRISPRko and (c) Replogle experiments. Rows indicate perturbations that have been hierarchically clustered into 4 groups. Columns are genes grouped into UPR pathway branch-specific genesets, as described in Adamson et al., 2016.

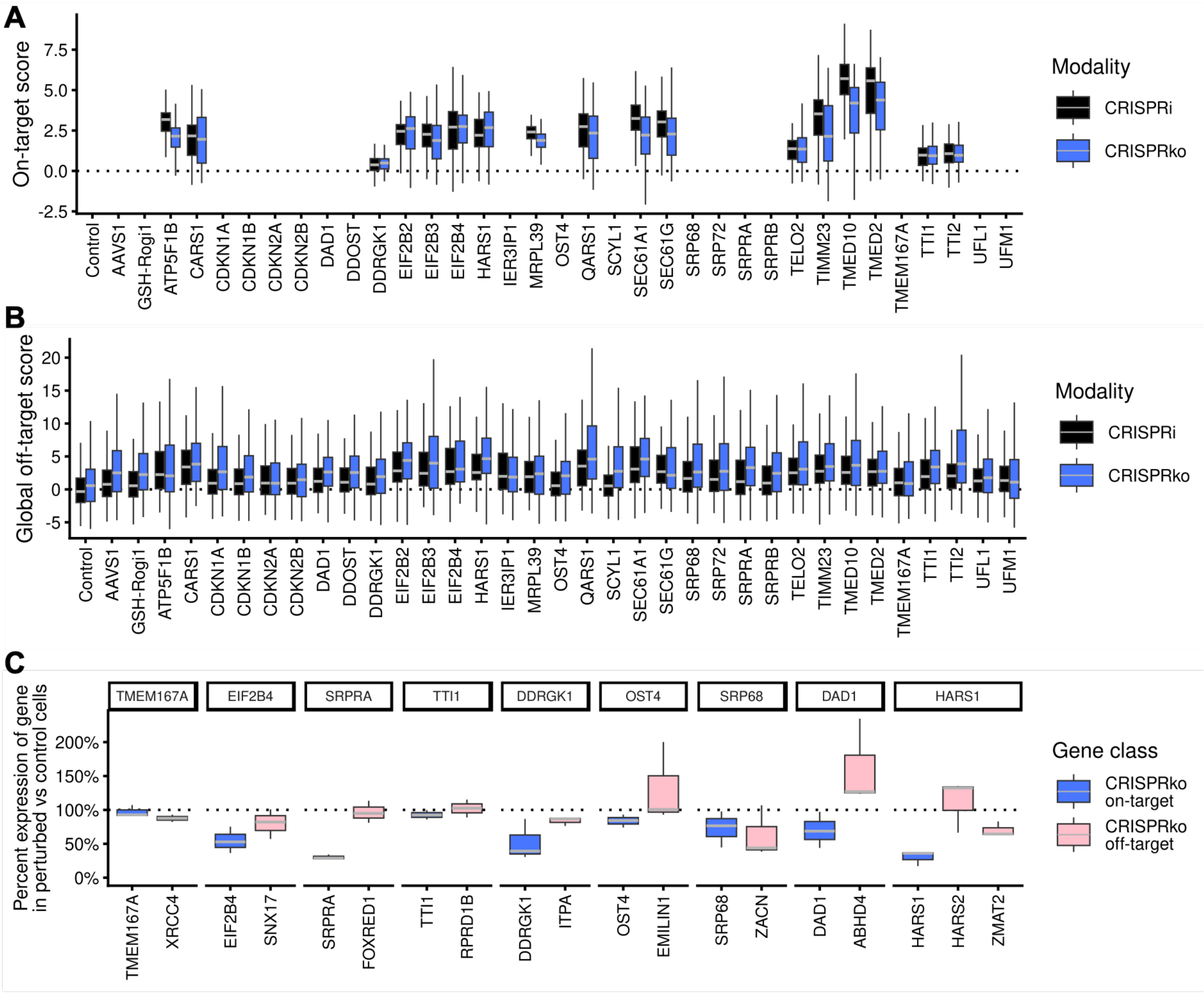


**Supplemental Figure 7.** a. For each perturbation, on-target perturbation scores were calculated by identifying shared differentially expressed genes between CRISPRi and CRISPRko (at least 5 were required), and using these to define a perturbation expression signature; every cell was projected onto its corresponding perturbation signature to yield an on-target score. For perturbations without sufficient differentially expressed genes, no on-target score could be calculated. Across all perturbations, higher on-target perturbation scores were observed with CRISPRi compared with CRISPRko p<2e-16. b. A global off-target score was calculated as the orthogonal residual between the total magnitude of expression changes and the on-target score (see methods). Across all perturbations, we observed that CRISPRko showed a higher off-target score compared with CRISPRi p<2e-16. c. Local off-target effects were assessed by identifying MANE Select transcripts with a transcriptional start site (TSS) within 10 kb of each CRISPRi target site. For perturbation-targets with a neighbouring TSS, the percentage gene expression of the perturbation target and neighbour in perturbed cells relative to expression of the same genes in non-targeting control (NTC) cells is shown for CRISPRko. No consistent repression of sgRNA target or neighbouring genes was observed.

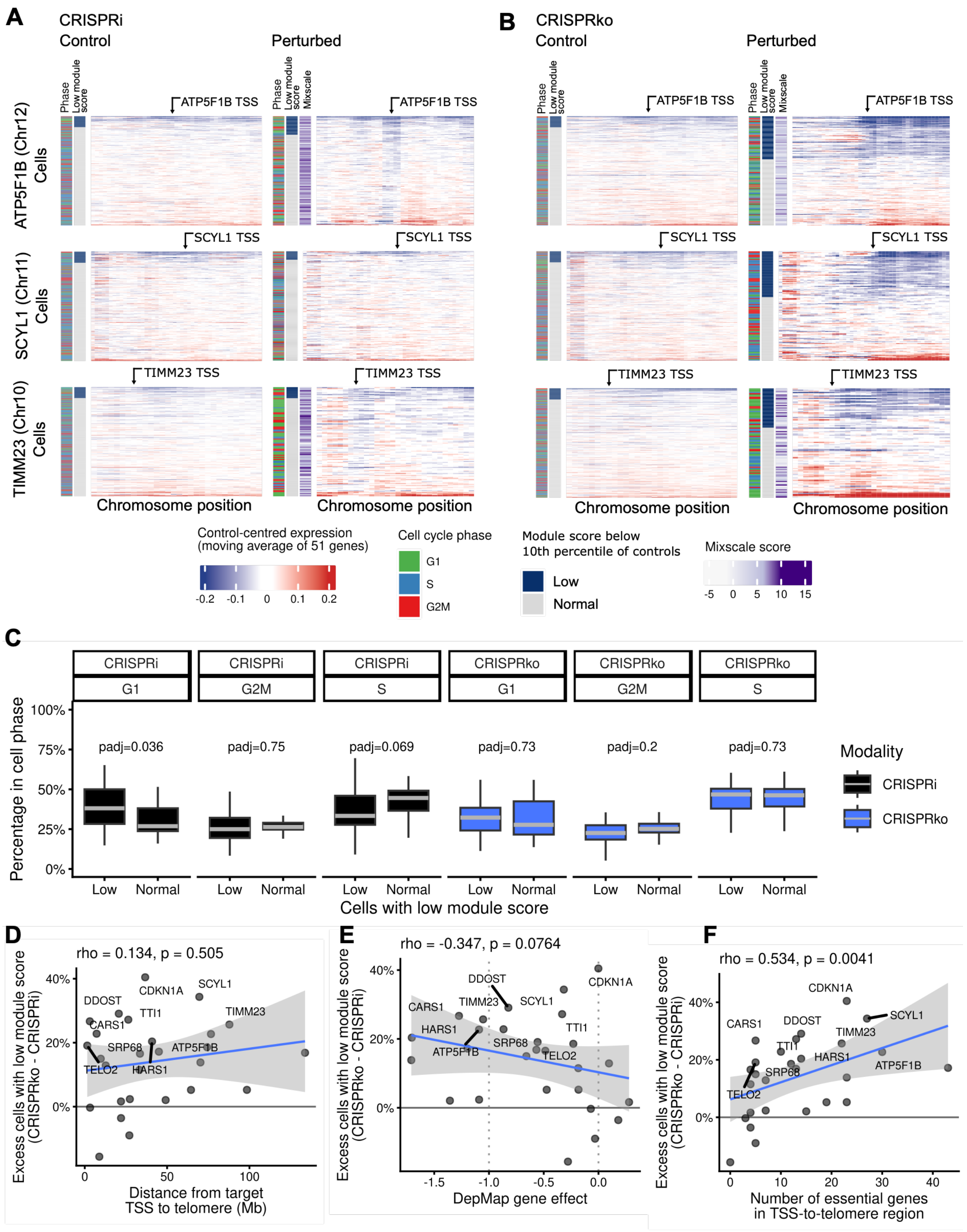


**Supplemental Figure 8.** a.and b. Three perturbations with evidence of telomeric loss of expression were selected as case studies to assess telomeric expression loss in further detail - ATP5F1B, SCYL1 and TIMM23. For each perturbation, a control-centred expression matrix was calculated. MANE select transcripts were ordered along the perturbation target chromosome, and a moving average (51 genes) of the control-centred expression was calculated. Each heatmap shows the moving average control-centred expression mapped to colour, for every cell (rows of heatmap) along the perturbation-target chromosome position (columns), with the perturbation-target TSS position marked by an arrow. Cells are ordered by their gene module score (Figure 2C). c. Percentage of cells per predicted cell cycle phase for each modality, perturbation and low/normal telomeric score classification (Figure 2C). Only CRIPSRi G1 phase showed a significant difference between low/normal telomeric classification ($padj < 0.05$). d. For every perturbation, the difference between CRISPRko and CRISPRi in the percent of cells with a low module score was calculated, resulting in a score for the excess percent of cells with a low module score in CRISPRko 'excess CRISPRko repression'. The distance between perturbation target sites and the end of chromosomes was plotted against this. There was not a significant correlation between distance to telomere and excess CRISPRko repression ($p > 0.05$). e. The DepMap gene effect for K-562 cells is plotted against the excess CRISPRko repression score; there was not a significant correlation with DepMap gene effect ($p > 0.05$). f. The number of essential genes (DepMap gene effect in K-562 below -1) found in the target TSS-to-telomeric region is plotted against the excess CRISPRko repression score. There was a significant correlation between number of essential genes and excess CRISPRko repression ($p < 0.01$).
